# Amygdala input differentially influences prefrontal local field potential and single neuron encoding of reward-based decisions

**DOI:** 10.1101/129221

**Authors:** Frederic M. Stoll, Clayton P. Mosher, Sarita Tamang, Elisabeth A. Murray, Peter H. Rudebeck

**Affiliations:** Icahn School of Medicine at Mount Sinai, One Gustave L. Levy Place, New York, NY, 10029, USA; Section on the Neurobiology of Learning and Memory, Laboratory of Neuropsychology, Building 49, Suite 1B80, 49 Convent Drive, Bethesda, Maryland 20892, USA

**Author notes:** **CORRESPONDENCE SHOULD BE ADDRESSED TO:** Dr. Peter H. Rudebeck Icahn School of Medicine at Mount Sinai One Gustave L. Levy Place New York, NY, 10029, USA.

**Keywords:** Amygdala, reward, affect, emotion, orbitofrontal cortex, local field potential, medial prefrontal cortex, decision making, choices, valuation, outcome

## Abstract

Reward-guided behaviors require functional interaction between amygdala, orbital (OFC), and medial (MFC) divisions of prefrontal cortex, but the neural mechanisms underlying these interactions are unclear. Here, we used a decoding approach to analyze local field potentials (LFPs) recorded from OFC and MFC of monkeys engaged in a stimulus-choice task, before and after excitotoxic amygdala lesions. Whereas OFC LFP responses were strongly modulated by the amount of reward associated with each stimulus, MFC responses best represented which stimulus subjects decided to choose. This was counter to what we observed in the level of single neurons where their activity was closely associated with the value of the stimuli presented on each trial. After lesions of the amygdala, stimulus-reward value and choice encoding were reduced in OFC and MFC, respectively. However, while the lesion-induced decrease in OFC LFP encoding of stimulus-reward value mirrored changes in single neuron activity, reduced choice encoding in MFC LFPs was distinct from changes in single neuron activity. Thus, LFPs and single neurons represent different information required for decision-making in OFC and MFC. At the circuit-level, amygdala input to these two areas play a distinct role in stimulus-reward encoding in OFC and choice encoding in MFC.

## SIGNIFICANCE STATEMENT

Dynamic interaction between amygdala, orbitofrontal (OFC) and medial frontal cortex (MFC) is required for adaptive foraging. To determine the nature of these neural mechanisms, we compared single neuron and local field potential responses (LFPs) in monkeys making reward-guided choices both before and after amygdala lesions. LFP responses in OFC best represented stimulus-reward values available on each trial, whereas MFC LFP responses were closely associated with monkeys’ choices. By contrast, single neurons, in both areas primarily encoded stimulus-reward value. Removing amygdala input to OFC and MFC heightened these differences between encoding of task variable by LFPs and single neurons. Thus single neurons and LFPs in frontal cortex represent different aspects of decision-making and are differentially influenced by the amygdala.

## INTRODUCTION

Interaction between the prefrontal cortex (PFC) and limbic system is required for normal patterns of affective behavior and cognition. In particular, reward-guided behaviors require functional interaction between the amygdala and the orbital and medial divisions of the PFC (OFC and MFC, respectively). For instance, lesions that disconnect the OFC and amygdala are associated with impairments in updating the value of rewards (Baxter et al., 2000). Similarly, disconnection of the MFC and amygdala leads to deficits in correctly weighting the costs and benefits of different courses of action (Floresco and Ghods-Sharifi, 2007) as well as emotional responding (Felix-Ortiz et al., 2016). Disruption of functional interaction between the PFC and limbic system, most notably the amygdala, is also associated with a host of psychiatric disorders (Pezawas et al., 2005; Almeida et al., 2009; Dutta et al., 2014). Determining how these brain areas interact at the neural level when one of the nodes of the network is dysfunctional or damaged is therefore a key first step to understanding circuit-level interactions.

We previously showed that in monkeys, bilateral excitotoxic lesions of the amygdala attenuate reward-value signals of individual neurons recorded from OFC, but not MFC (Rudebeck et al., 2013). The response properties of single neurons, however, only reflect the local processing and output of an area (Einevoll et al., 2013). A more complete understanding of this network might be informed by considering population-level activity and the inputs to an area, which can be studied using local field potentials (LFPs). Indeed there is evidence that there are differences in the types of information encoded by single neurons and LFPs (Kreiman et al., 2006). For example, during a working memory task where object locations and features had to be held online, differences emerged between the information encoded in spike trains and LFPs in lateral frontal cortex (Lara and Wallis, 2014). In the context of the present data, we previously reported that stimulus-reward encoding in OFC was independent of concurrently presented options, suggesting that the activity of single neurons was not encoding the monkeys’ choices (Rudebeck et al., 2013). However, it is possible that choices might be being encoded at the level of LFPs instead of the activity of single neurons in OFC and MFC.

To better determine the dynamics of reward-value and choice coding in the amygdala-MFC-OFC network, we analyzed LFPs recorded from the OFC and MFC in three monkeys engaged in a stimulus-choice task. Recordings were made both before and after excitotoxic lesions of the amygdala. Specifically we looked for signals that were associated with encoding stimulus-reward associations and monkeys’ choices and compared this to encoding by single neurons. Here we report that prior to lesions of the amygdala we could, mirroring the spike data, decode stimulus-reward values in OFC to a greater extent than MFC. By contrast, the choice that would ultimately be taken was more strongly encoded in MFC than OFC. Removing amygdala input led to reduced signals in OFC and MFC related to stimulus values and choices, respectively.

## METHODS

### Subjects

Three adult male rhesus macaques (*Macaca mulatta*), monkeys H, N and V, served as subjects; they weighed 8.5, 8.0 and 8.4 kg, respectively, at the beginning of training. Animals were pair housed when possible, kept on a 12-h light dark cycle and had access to food 24 hours a day. Throughout training and testing each monkey’s access to water was controlled for 6 days per week. All procedures were reviewed and approved by the NIMH Animal Care and Use Committee.

### Apparatus

Monkeys were trained to perform a two-choice visually guided task for fluid reward. All trial events and timing were controlled using the open source NIMH Cortex program (ftp://helix.nih.gov/lsn/cortex/). Eye position and pupil size were monitored and acquired at 60 frames per second with an infrared occulometer (Arrington Research, Scottsdale, AZ).

During training and testing monkeys sat in a primate chair with their heads restrained. Directly in front of the chair, three buttons were spaced horizontally 7 cm apart (center to center). These buttons had embedded infrared sensors to detect contact.

### Task and behavior

Three monkeys were trained to perform a choice task for fluid rewards. On each trial, monkeys had to press and hold a central button and then fixate a central light spot for 0.5–1.5 s (Fig. 1A). Two visual stimuli, associated with different amounts of fluid reward, were then sequentially presented. The onset of the second stimulus (S2) followed the onset of the first (S1) by 1.0 s and, by random selection, one stimulus appeared to the left of the central spot and one appeared to the right. We presented the two stimuli for choice sequentially in an attempt to separate the valuation process of the individual items. Stimuli were randomly selected from a pool of ten stimuli (Fig. 1A). Monkeys had learned that each of the stimuli was associated with a fixed amount of fluid — 0.8, 0.4, 0.2, 0.1 or 0 ml of water — two stimuli for each quantity. A total of 14 pairs were tested (S1/S2 values): 0/0.1 ml, 0/0.2 ml, 0.1/0 ml, 0.1/0.2 ml, 0.1/0.4 ml, 0.2/0 ml, 0.2/0.1 ml, 0.2/0.4 ml, 0.2/0.8 ml, 0.4/0.1 ml, 0.4/0.2 ml, 0.4/0.8 ml, 0.8/0.2 ml, and 0.8/0.4 ml. All sessions also included pairs with stimuli associated with the same reward values for S1 and S2 (i.e. 0.1/0.1 ml, 0.2/0.2 ml and 0.4/0.4 ml) on 10% of the trials. Given the limited number of these trials, they were not included in the present analyses. After a variable delay of 0.0–1.5 s, the central spot brightened as a “go” signal, and the monkeys could then choose between the two stimuli by reaching to the left or right response button. The amount of fluid reward corresponding to the chosen stimulus was delivered 0.5 s later.

**Figure 1.**
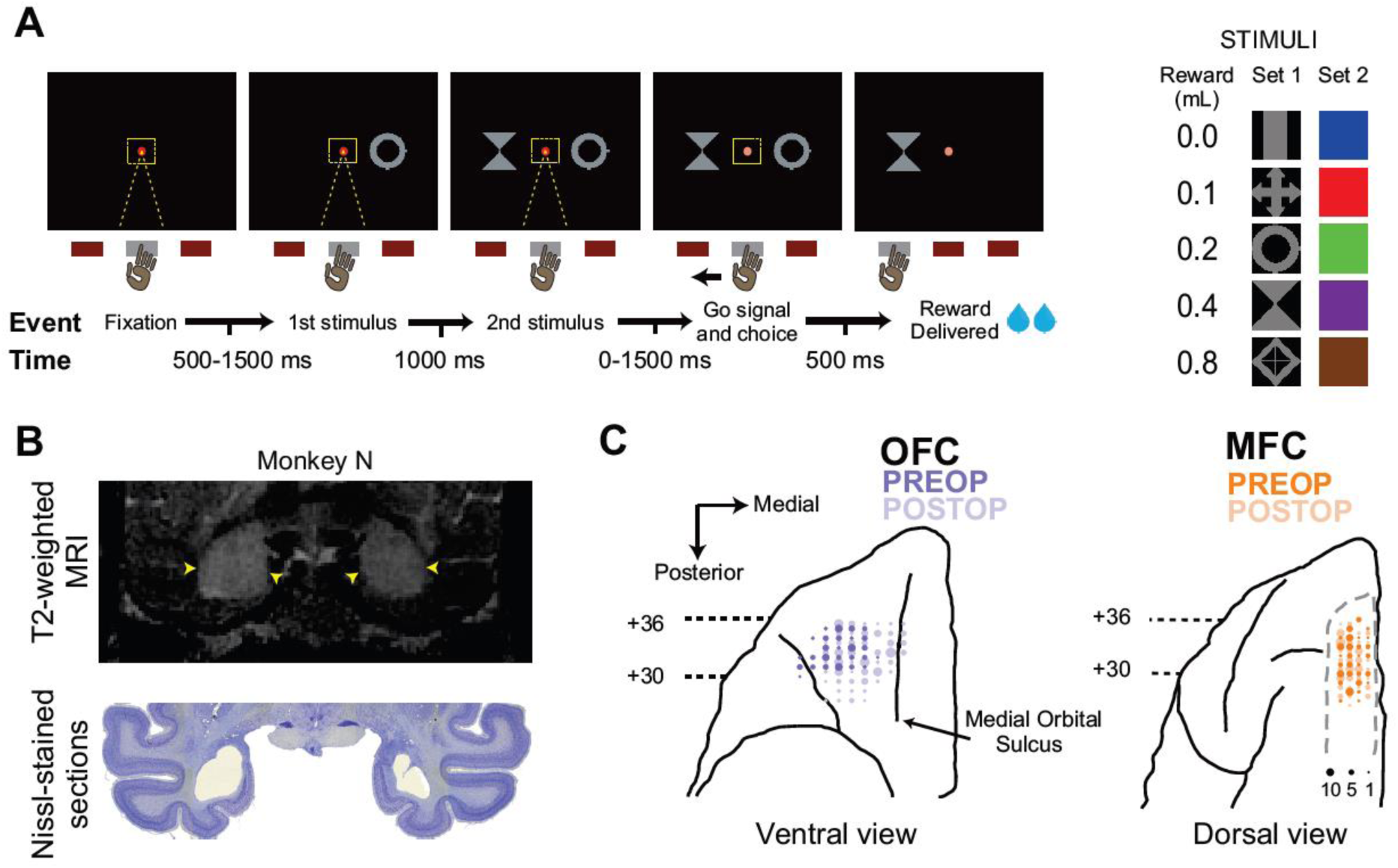
Two-choice reward-guided task and recording positions. (**a**) On each trial, two stimuli were sequentially presented on the right and left side of the screen (randomized from trial to trial) while monkeys maintained central fixation. After a random delay, the central fixation spot changed color, and monkeys were allowed to select with their hand the stimulus of their choice. The reward amount associated with the selected stimulus was then delivered. Two sets of stimuli were used (shapes or color), each containing 5 different stimuli associated with 0, 0.1, 0.2, 0.4 or 0.8 ml of water. (**b**) T2-weighted MRI (top) and post-mortem Nissl-stained section (bottom) illustrating the extent of the bilateral excitotoxic amygdala lesion performed in one representative monkey (see Rudebeck et al., 2013 for a complete description). (**c**) Preoperative (dark colors) and postoperative (light colors) recording locations in both the OFC (left) and the dorsal bank of the MFC (right). Dot sizes indicate the number of recordings for each site. Antero-posterior values indicate distance in mm relative to the ear bars.

### Surgical procedures, neural recordings, imaging and histological reconstruction

For detailed information on surgical procedures, see Rudebeck et al. (2013). In brief, each monkey was implanted with a titanium head restraint device and then, in a separate surgery, a plastic recording chamber was placed over the exposed dura mater of the left frontal lobe. After the preoperative recordings were completed, MRI-guided bilateral excitotoxic lesions of the amygdala were made in each monkey.

Potentials from single neurons and local field potentials were recorded with tungsten microelectrodes (FHC, Inc. or Alpha Omega, 0.5-1.5 MΩ at 1 KHz) advanced by an 8-channel micromanipulator (NAN instruments, Nazareth, Israel) attached to the recording chamber. Spikes from putative single neurons were isolated online using a Plexon Multichannel Acquisition Processor and later verified with Plexon OffLine Sorter on the basis of principal-component analysis, visually differentiated waveforms and interspike intervals. Neurons were isolated before monkeys were engaged in any task. Other than the quality of isolation, there were no selection criteria for neurons. Local field potentials were recorded using the same system and digitized at 1kHz. Recordings were referenced on guide tubes containing the electrodes and in contact with the dura.

OFC recordings were made on the ventral surface of the frontal lobe between the lateral and medial orbital sulci, roughly corresponding to Walker’s areas 11 and 13. All OFC recordings were between +27 and +38 mm anterior to the interaural plane. Recording locations in MFC were primarily in the dorsal bank of the cingulate sulcus (areas 9 and 24), although some sites were in the ventral part of the fundus of the cingulate sulcus. MFC recordings were made between the anterior tip of the cingulate sulcus (approximately +38 mm) and +24 mm.

Both before and after lesions of the amygdala, recordings were made in overlapping regions in each of the three monkeys (Fig. 1C). Recording sites were verified by T1-weighted MRI imaging of electrodes after recording sessions and by placing electrolytic marking lesions (15 μA direct current for 25 seconds, electrode positive) at selected locations in OFC after recordings had been completed (see Rudebeck et al., 2013). At the conclusion of the study, monkeys were deeply anesthetized and transcardially perfused with saline (0.9%) followed by formalin. The brains were removed, sectioned in the coronal plane, Nissl-stained and mounted onto glass slides for visual inspection.

The extent and location of the amygdala lesions was assessed using T2-weighted MRI conducted within one week of each surgery (e.g. Fig. 1B, top row). Lesion volume was then confirmed from histology (e.g. Fig. 1B, bottom row). The locations and extents of the lesions were largely as intended. There was near complete cell loss in all nuclei in the amygdaloid complex (mean = 95.5%). Inadvertent damage was evident in the entorhinal and perirhinal cortex, portions of the ventral claustrum, and anterior hippocampus (see Rudebeck et al., 2013). Importantly, with the possible exception of the entorhinal cortex, this unintended damage was slight (e.g., extending less than 2 mm in antero-posterior extent) and asymmetric between the hemispheres. Finally, one monkey (Monkey N) sustained an infarction in the dorsal striatum, bilaterally. Overall, damage in all three monkeys consistently centered on the amygdala, bilaterally.

### Electrophysiological data processing

Data analysis was performed offline using the FieldTrip toolbox (Oostenveld et al., 2011) and custom Matlab scripts (Matlab, MathWork Inc.).

Preoperatively, a total of 234 and 155 LFPs were recorded from 3 monkeys in the OFC and MFC respectively. Following the bilateral amygdala lesions, we recorded 324 and 204 more LFPs in the OFC and MFC, respectively. To avoid biasing our results, we required a minimum of 5 trials for each possible pair of S1 and S2 values, and we did not include trials in which there was no delay between S2 presentation and the Go signal (i.e., delay needed to be at least 500 ms or more). A few recordings did not meet the trial number requirement and were therefore excluded from further analyses (PreOp: OFC=12/234, MFC=11/155 sites; PostOp: OFC=26/324, MFC=20/204 sites).

#### Pre-processing

Here, our analyses focused specifically on the event-related potentials (ERPs) evoked by stimuli presentation. We did not report data on possible oscillatory activities as no clear modulations have been observed apart from the direct ERP signature in the low frequency spectrum (data not shown). For each recorded site, the LFP signal was first band-pass filtered from 1 to 30 Hz and then aligned around S1 presentation (from -3 to +3 s). We then normalized the LFP signal for each individual trial relative to a baseline period (-0.6 to -0.1 s before S1 onset), and derived a z-score. Finally, we sub-sampled our dataset using a sliding window of 50 ms stepped in 10 ms increments.

#### ERPs latency and amplitude

The latency of the different ERP components was extracted for each trial by detecting peaks with amplitude greater than 0.3 sd and with a minimal distance restriction between 2 consecutive peaks of 75 ms (using the Matlab function *findpeaks.m*). Latencies for negative components were considered only if they fell between 200 and 350 ms relative to stimulus onset, whilst latencies for the later positive components needed to be between 300 and 550 ms. These time windows were defined based on the visual inspection of all the detected latencies. We then used Kruskal-Wallis (KW) tests to assess differences between conditions.

Finally, differences in ERP amplitudes between areas (OFC vs. MFC) or relative to amygdala lesions (PreOp vs. PostOp) were also extracted using KW tests at each time bin.

#### ANOVA on individual LFPs

To assess how the different factors of our task modulated the amplitude of the trial-by-trial ERP signals, we first fitted a sliding hierarchical ANOVA model to the normalized ERP activity of each recorded LFP. Our model included factors of S1 reward value (five levels), S2 reward value (five levels), S1 identity (two levels), S2 identity (two levels) and S1 presentation side (two levels). S1 and S2 identity factors were nested within S1 and S2 reward value, respectively. P-values extracted for each factor were subjected to a specific threshold to account for multiple testing over time (see *Statistical procedures* below). To complement our time-resolved observations, we also extracted the overall number of significant sites for 3 different periods: a reference period (REF: -1000 to 0 ms, relative to S1 onset), the S1 period (0 to 1000 ms) and S2 period (1000 to 2000 ms). Finally, to extract the latency of stimulus value or stimulus side encoding, we detected the first significant time bin during the time period considered (S1 or S2 periods). As before, differences in latencies were evaluated using KW tests.

#### Statistical procedures applied to individual LFPs analyses

Individual sites were considered as significantly encoding a task factor if they discriminated that factor for 6 consecutive bins (covering a time period of 100ms) with a threshold of p<0.01. This threshold was applied to all time-resolved analyses (e.g. hierarchical ANOVA and KW test on ERP amplitude over time).

We also assessed statistical significance by computing two-tailed Chi-square tests with Yates’ correction when testing for differences in the proportion of sites encoding a given factor, either when comparing areas (OFC vs. MFC) or periods (PreOp vs. PostOp).

#### Relationship to single neuron recordings

We previously reported the encoding properties of the individual neurons recorded in this task (Rudebeck et al., 2013; PreOp: OFC=280; MFC=233; PostOp: OFC=317; MFC=237 neurons). Here, we were able to assess whether there was a relationship between neurons and ERPs as the same analyses were applied with the two datasets. In particular, we tested whether neurons and ERPs recorded simultaneously on the same electrode (i.e., at a similar site) were modulated by the different factors in a similar manner. To do this, we extracted the proportion of sites where, based on the hierarchical ANOVAs, both neurons and ERPs showed significant encoding of S1 or S2 values. To test if these proportions were significant (e.g., whether neurons recorded on an electrode showing significant ERP modulations were more likely, or not, to also encode the same factor), we used permutation testing, by shuffling the labels assigned to the different neurons (significant or not) 1000 times. This procedure enabled us to take into account the relative number of both significant LFP sites and neurons, and therefore avoid any confounds.

We also looked at the correlation between the variance explained by the S1 or S2 values from the hierarchical ANOVAs in recordings where both neurons and ERPs (recorded on the same electrode) showed significant modulations.

#### Population decoding of choices and stimulus reward values

We applied multiple linear regressions to decode information from population ERP activity vectors in both regions. This method assesses the capability of a linear readout to extract a given response variable (e.g., choosing S1 or S2) from trial-by-trial ERP responses of the whole population. In this procedure, a Tikhonov regularization procedure was used to minimize the sum of squared errors and thus avoid overfitting by placing constraints on regression coefficients.

To extract an accurate estimate of the classifiers’ performance, we included only sites with 3 repetitions of each of the 14 possible S1/S2 pairs. We also only included the same number of predictors (i.e., recording sites) for both areas, as well as for pre-and post-operative recordings. This was done so that we could directly compare the strength of coding between the different recording populations, as more predictors might spuriously increase the accuracy (see for example, Astrand et al., 2014). Applying these criteria meant that the datasets used to extract choice-predictive activity contained the ERP signals recorded at 47 randomly selected sites during 42 randomly selected trials. Our training set contained 2 instances of each possible pair (i.e., 28 trials), and our testing set contained the remaining third of the data (14 trials). It is important to note that classifiers did not have any information relative to the S1 or S2 reward values, nor did it have information regarding S1/S2 pair identities. To determine the regularization parameter, we further subdivided the training partition to perform a 5-fold cross-validation procedure. The sum of squared errors (SSE) across the five folds was computed for each regularization parameter tested. The value with the lowest SSE was then selected and used to train the classifier on the whole training partition. We then tested the classifier on the remaining testing partition, which contained the 14 trials. This meant that the testing partition was never used during the optimization and/or training of the classifier. Given our under-sampling procedure, an unbiased readout of performance was extracted by randomly selecting trials and performing all computations 1000 times, which generated a vector of 1000 estimated choices for each of the 14 S1/S2 pairs. We then used the average choice binary output over these 1000 computations. Classifiers were trained and tested at each time bin.

Chance levels and statistical significance were defined using a permutation approach. Specifically, we randomly permuted monkeys’ choices 1000 times without removing the relationship between ERP signal and the different S1/S2 pairs and conducted the same decoding approach described above. Importantly, the temporal structure of the ERP signal also remained unaffected by the permutation procedure. All subsequent computations were done in a similar manner to that described above. Finally, classifiers’ performances were averaged across the 14 S1/S2 pairs for each permutation and used to assess statistical significance.

The same analysis methods were also applied to the recordings of single neuron activity (OFC and MFC neurons, both before and after lesions). Here we used the activity of a subsample of 47 randomly selected neurons for each population of neurons from OFC and MFC to predict monkeys’ choices. Just as we had done for the ERP analyses, described above, we first used the average firing rate for 50 ms bins each 10 ms step. With such time averaging, we were almost unable to decode choices from either OFC (PreOp=1/14 and PostOp=3/14 significantly decoded S1/S2 pairs) or MFC (PreOp=2/14 and PostOp=0/14). However, it is unclear if this null result was due to a limitation inherent to the nature of the signal (i.e., spiking activity is a point-process, as opposed to the continuous ERP signal) or a true absence of encoding. Our objective being the comparison between neuronal and ERP populations, we therefore reported the decoding performance using longer bins of 200 ms for neuronal populations, a common window size used to analyze neuronal activity (e.g. Rudebeck et al., 2013; Lara and Wallis, 2014; Stoll et al., 2016).

We report the results of our time-resolved approach, but also after averaging 2 time periods (*t1*, from 300 to 400 ms; and *t2*, 1250 to 1350 ms, relative to S1 onset). These time windows were defined to match the ERPs components and based on the overall decoding performance. Statistical estimates for these time windows were extracted by averaging the results of the 1000 permutations over time in a similar manner. Differences between recorded populations were assessed using Chi-square tests with Yates’ corrections. We used a threshold of p<0.05 after correcting for multiple tests.

To further compare the decoding performance for the different conditions, we fitted a mixed-effect logistic regression on the output of the classifiers (average number of S1 choices out of the 1000 permutations during time bin *t2*). The full model included fixed-effect for all three categorical fixed-effect factors (Type: neurons vs. LFPs; Area: OFC vs. MFC, Surgery: PreOp vs. PostOp) and all interactions. Also, S1/S2 pairs were dummy-coded and included as a random-effect factor, allowing changes in intercept (i.e., choosing more S1 or S2 depending on their respective values). Thus, this model compared the overall performance of the classifiers independently of the S1/S2 pair considered. Model coefficients were derived by maximum likelihood estimation using Laplace approximation. We report the output from the full model in the results given that a similar model without the 3-way interaction (Type x Area x Surgery) was significantly less adapted to fit our dataset (Log-Likelihood test, LR=37.6, p=8.7e-10). We also validated our model by ensuring that normalized residuals plotted against fitted values and factors did not show inhomogeneity.

Finally, we investigated whether it was also possible to extract stimulus-reward values using population decoding methods. Here, we applied support vector machine (SVM) algorithms with Gaussian kernel on the ERP signals recorded from 47 randomly selected channels. This procedure was performed to extract S1 and S2 values using the average ERP amplitude during time bins t1 and t2, respectively, for each trial and each predictor (recording sites). We used the ERP activity of 10 randomly selected trials for each S1 or S2 values (for a total of 50 trials). Readout performances were extracted by randomly selecting trials and performing all computations 100 times. We then averaged the decoding performance and compared it with a set of 1000 randomly-generated permutations.

#### Experimental Design and Statistical Analysis

Three adult male rhesus macaques (*Macaca mulatta*) were used in this study. Recordings in OFC and MFC were made in each monkey before and after amydgala lesions. This means that each monkey served as its own control. Statistical comparisons (OFC vs. MFC, PreOp vs. PostOp, ERPs vs. single neurons) were performed at the level of the population as well as for each subject when possible (see above). For count-based data statistics we used chi-square tests and where appropriate, ANOVAs, Kruskal-Wallis or permutation tests for continuous data.

## RESULTS

### Task and Behavioral performance

Three monkeys were trained to perform a two-choice reward-guided task for fluid rewards (see Methods, Fig. 1A). On each trial, monkeys had to choose one of two visual stimuli, associated with different amounts of reward, that were sequentially presented. Stimuli were randomly selected from a total of ten stimuli, each one associated with a fixed amount of fluid (0, 0.1, 0.2, 0.4 or 0.8 ml, two stimuli for each quantity, Fig. 1A).

Behavioral performance during the task has been described in detail elsewhere (Rudebeck, et al., 2013). In brief, each monkey chose the stimulus associated with the greatest amount of reward on nearly every trial (>95%). Bilateral excitotoxic lesions of the amygdala did not alter monkeys’ performances; monkeys continued to select the stimulus associated with the greatest amount of reward on more than 95% of trials. Although it might seem counterintuitive, the task was specifically designed, based on prior work (Izquierdo and Murray, 2007), to ensure that performance would not be affected by the lesions. This aids the interpretation of the results; if there had been a behavioral deficit postoperatively it would be difficult to interpret any postoperative changes in neural activity, as effects could be due to either the lesion or the change in behavior. In addition, we confirmed that the lesions were effective in a separate task that required the learning of new stimulus-reward associations (Rudebeck et al., 2017).

Choice response times, defined as the amount of time between the go signal being delivered and the monkey lifting its hand to make a movement, were modulated by the amount of reward that the monkeys would receive for making a particular choice (p<0.01, see Rudebeck et al., 2013). Lesions of the amygdala did not consistently alter the effect of value on monkeys’ choice latencies (p>0.3).

### Encoding of stimulus value in the ERP

While monkeys performed the task, we recorded both single neurons and local field potentials in the OFC and MFC. Here, we report the analysis of a total of 366 LFPs (OFC=222 sites, MFC=144 sites) recorded from the 3 monkeys before bilateral excitotoxic lesions of the amygdala (monkey N: 81 OFC, 77 MFC; monkey H: 20 OFC, 56 MFC; monkey V: 121 OFC, 11 MFC). The presentation of each stimulus (S1 or S2) was associated with a strong ERP response at both OFC and MFC recording sites (Fig. 2A,D). The early visual responses induced by the presentation of S1 were followed by two main components: a negativity peaking around 250 ms (average ± std, OFC = 274.4 ± 28 ms; MFC = 259.4 ± 21 ms) followed by a late positivity around 400 ms (OFC = 433 ± 52 ms; MFC = 411.6 ± 48 ms). Both the early negativity and late positivity were significantly earlier in the MFC compared to the OFC (KW test; negativity: H=21.04, p=4.49e-6; positivity: H=15.46, p=8.43e-5). The presentation of S2 elicited similar ERP responses to those following S1.

**Figure 2.**
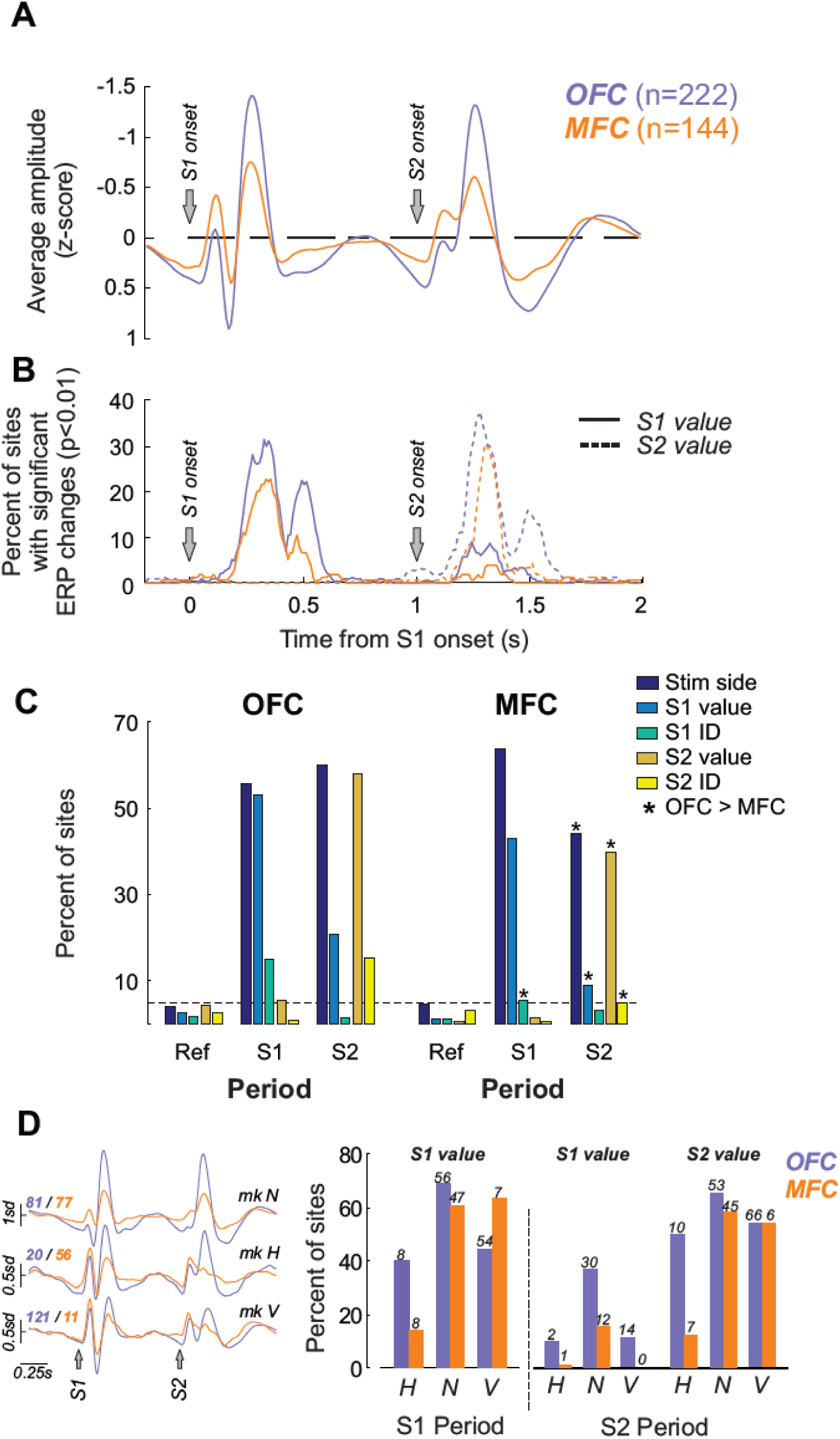
Stimulus-reward modulation of ERPs in the OFC and MFC. (**a**) Grand average normalized ERP responses induced by the presentation of S1 and S2 in the OFC (purple) and MFC (orange). Black lines indicate significant differences in power between OFC and MFC (KW test, p<0.01). (**b**) Time-resolved percentage of significant sites (hierarchical ANOVA thresholded at p<0.01, see Methods) encoding either S1 (solid lines) or S2 values (dashed lines) in both OFC and MFC. (**c**) Percentage of sites in the OFC (left) and MFC (right) showing a significant effect of one of the factors in the hierarchical ANOVA for the 3 different time periods (Ref: -1 to 0 s; S1: 0 to 1 s; S2: 1 to 2 s). Stars indicate a significant difference between OFC and MFC for the factor considered (Chi-square tests, p<0.05). (**d**) Individual monkeys’ normalized ERP responses (left) and percentage of sites in the OFC and MFC showing a significant effect of S1 and S2 values either during S1 or S2 periods (right). Numbers on top of bars indicate the numbers of significant sites. H, N and V represent the three monkeys.

To characterize the relationship between stimulus values and the different components of the ERP responses, we performed a sliding hierarchical ANOVA based on single-trial responses around the presentation of both S1 and S2 (see Methods). Stimulus value coding was found in both OFC and MFC ERPs, in particular at the time of the described early negativity and later positivity of the ERP responses (Fig. 2B). No difference in the latency of value coding between OFC and MFC was found for the encoding of S1 values (average ± std, OFC = 282.4 ± 102 ms; MFC = 275.9 ± 121 ms; KW test, H=0.03, p=0.8606). However, S2 value was encoded earlier in OFC compared to MFC (average ± std, OFC = 237.7 ± 106 ms; MFC = 278.7 ± 129 ms; KW test, H=8.47, p=0.0036).

To further quantify the contribution of OFC and MFC to stimulus reward-value coding, we looked at the proportion of sites encoding each factor during 3 time periods around stimulus presentation (Reference period: -1 to 0 s; S1 period: 0 to 1s; S2 period: 1 to 2 s, all relative to S1 onset, Fig. 2C). During the S1 period, the encoding of S1 values was observed at more OFC sites than MFC sites (OFC=118/222, 53.15%; MFC=62/144, 43.05%; **X**^2^=3.56, p=0.059). A similar pattern was seen during the S2 period (OFC=129/222, 58.1%; MFC=58/144, 40.2%; **X**^2^=11.11, p=0.0009). However, only monkeys H and N displayed a consistent difference between areas for S1 and S2 values (Fig. 2D). No differences were observed in monkey V, although this is likely due to the small number of recorded sites in the MFC (n=11). Finally, a small percentage of sites also encoded the value of S1 during the S2 period, and again this was higher in OFC (OFC=20.7% and MFC=9%, **X**^2^=7.988, p=0.0047, Fig. 2C). It should be noted that S1 remained on the monitor screen at the time of S2 presentation. The percentage of sites that coded S1 value during the S2 period was consistently higher in the OFC than the MFC of all three monkeys (Fig. 2D).

Recording sites showing an encoding of the value of S1 in the ERPs during S1 period were highly likely to encode the value of S2 during the S2 period in both the OFC (n=90/118, 76.3%) and the MFC (n=33/62, 53.2%). The encoding of both S1 and S2 values in similar sites was significantly greater in the OFC than the MFC (**X**^2^=8.939, p=0.0028). Qualitative inspection of the data did not reveal any topological differences related to stimulus value coding across the anatomical extent of either OFC or MFC.

Importantly, the encoding of reward values can also be extracted using population measures, by applying nonlinear Support Vector Machines (SVM) with a Gaussian kernel. Both OFC and MFC populations discriminated S1 values when the stimulus was presented (OFC: decoding rate ± std = 60.9±11.1 %, p=0.001; MFC: decoding rate = 49.4±10.4 %, p=0.001; average permutation chance level being 20%), with higher decoding performance in the former (KW test, H=45.92, p=1.23e-11). A significant discrimination of S2 values was also observed (OFC: 58±10.5 %, p=0.001; MFC: 47±9.5 %, p=0.001), again with better performance in the OFC population than the MFC one (KW test, H=47.2, p=6.35e-12).

Additional analyses revealed that the side on which the stimulus was presented (randomized from trial to trial, see Methods) explained a sizable proportion of the ERP responses (Fig. 2C). While there was no apparent difference between OFC and MFC during S1 presentation (OFC=55.8% and MFC=63.8%, **X**^2^=2.33, p=0.13), we found significant differences during S2 presentation (OFC=59.9% and MFC=44.4%, **X**^2^=8.40, p=0.0037). However, large discrepancies between monkeys, both during S1 (monkey H: OFC=9/20, MFC=21/56; monkey N: OFC=76/81, MFC=71/77; monkey V: OFC=39/121, MFC=0/11) and S2 period (monkey H: OFC=6/20, MFC=12/56; monkey N: OFC=75/81, MFC=50/77; monkey V: OFC=52/121, MFC=2/11), mean that this result should be treated with caution. Contrary to the encoding of stimulus value, the modulation of the ERP by the stimulus side was almost exclusively observed during the initial ERP responses (stimulus side discrimination peaked at 225 ms and 215 ms after S1 onset for OFC and MFC respectively).

We also found that the encoding of the identity of S1 or S2, either color or shape stimuli, was only apparent in the OFC (S1=33/222, 14.8%; S2=34/222, 15.3%). Only ∼5% of sites in MFC signaled stimulus identity (S1=8/144, 5.5%; S2=7/144, 4.8%). This was lower than in OFC (**X**^2^>7.6, p<0.0058) and also no different to chance levels.

In summary, these analyses of the ERPs from OFC and MFC reveal that: 1) OFC exhibited more prevalent and reliable encoding of stimulus-reward values compared to MFC; 2) stimulus location modulated the early ERP component in both OFC and MFC; 3) sites in the OFC and MFC encoding of the value of S1 during the S1 period were highly likely to also encode the value of S2 during the S2 period, and 4) stimulus identity encoding was only evident in OFC, not MFC.

### Comparison of ERPs and single-unit encoding of stimulus value

As reported by Rudebeck et al (2013), single units recorded in both OFC and MFC during performance of the task encoded stimulus values, i.e., the anticipated value of the reward outcome associated with a stimulus. Here, we investigated whether ERPs encoding stimulus values were more likely to be recorded on electrodes where the spiking activity of single neurons also encoded stimulus values. In total, 83.3% and 87.5% of analyzed LFP sites in OFC and MFC respectively (OFC=185/222 and MFC=126/144 sites) contained at least one well-isolated and simultaneously recorded single neuron. When we considered only these sites, similar effects to the ones previously described were found for the percentage of sites showing ERP encoding of S1 and S2 value (S1 value during S1 period, OFC=50.8% and MFC=42.06%; and S2 value during S2 period, OFC=58.38% and MFC=39.68%). We then looked at whether single neurons simultaneously recorded at these sites also encoded stimulus values. We found that single neurons encoding either the value of S1 (OFC=73/185 and MFC=36/126 neurons) or S2 (OFC=58/185 and MFC=22/126 neurons) were no more likely to be recorded at sites where ERPs also represented stimulus values. This was true for both for S1 (OFC=37/73, 50.7% and MFC=17/36, 47.2%) and S2 (OFC=37/58, 63.8% and MFC=9/22, 40.9%). None of these proportions were greater than expected by chance (permutation tests p>0.18). We found a similar pattern of results when we compared the explained variance related to S1 and S2 values encoded by either single neurons or ERPs (S1 value: OFC, r=0.08, p=0.62; MFC, r=0.15, p=0.57; S2 value: OFC, r=0.25, p=0.13; MFC, r=0.32, p=0.44). Therefore, our findings demonstrate that there is no direct relationship between the encoding of value in single neurons and ERPs simultaneously recorded at the same site in this task.

### Population encoding of choices during stimuli presentation

To further explore how reward-value signals in OFC and MFC might contribute to choice behavior, we applied multiple linear regressions to decode monkeys’ choices on each trial from ERP population activity vectors in both regions (see Methods and Fig. 3). Here when we refer to the choice, we mean the option, either S1 or S2 that the monkey will subsequently choose on each trial. Linear classifiers were trained on a subset of trials to discriminate the 2 categories: choosing S1 (=1) or choosing S2 (=-1). The training set contained 2 instances of each of the 14 possible S1-S2 pairs; the testing set contained 1 instance of each (total number of trials used was 48). This procedure was performed in order to retrieve *a posteriori* the classifier’s performance for each trial type without any bias in the number of their occurrences. Note that neither information concerning the identity of the different pairs, nor the value of S1 or S2, was given to the classifiers; only the chosen stimulus (S1 or S2) was used. It is also important to keep in mind that monkeys’ choices in this task are entirely based on the reward values associated with the different stimuli. Because monkeys nearly always chose the stimulus associated with the highest amount of reward, fully disentangling value and choice-related signals is beyond the scope of this study.

**Figure 3.**
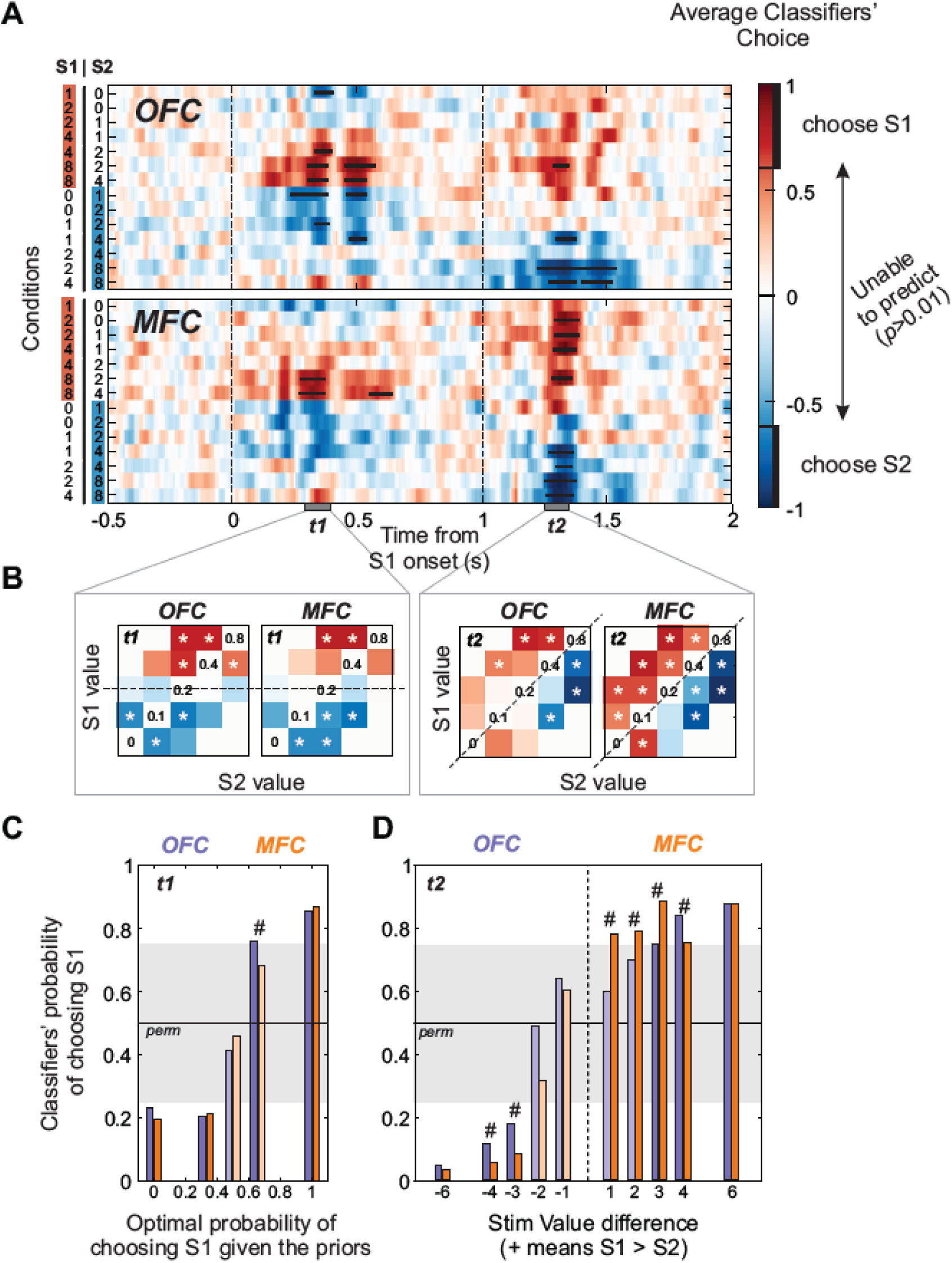
Population encoding of choices during stimuli presentation. (**a**) Average classifiers’ choice performance (red=S1 and blue=S2; a value of 1 or -1 means always choosing S1 or S2 respectively) for each individual pairs of stimuli (labeled as *Conditions*, y-axis) over time (x-axis). Black bars represents a significant statistical preference for choosing S1 or S2 based on permutation testing (threshold at p<0.01 for at least 6 consecutive time bins). (**b**) Time bin averages (for t1 and t2) of classifiers’ choices in each condition. Stars indicate significant decoding performance. (**c**) Average classifiers’ probability of choosing S1 during time bin t1 against the optimal probability of such choice given the S1 value and the task design (see Methods). (**d**) Average classifiers’ probability of choosing S1 during time bin t2 against stimuli value difference. In panel c and d, # indicates significant differences between OFC and MFC. Gray shading represent the noise level extracted from permutations. OFC=Purple; MFC=Orange bars. Dark and light colors show significant and non-significant probabilities respectively.

Decoding monkeys’ choices was studied here using ERP population activity vectors combining sites from all monkeys. Due to our trial number requirement, the total number of available recording sites for the different monkeys varied in both the OFC (monkey H=2, monkey N=51, monkey V=59 sites) and MFC (monkey H=8, monkey N=35, monkey V=3 sites) populations. As a result, most of the sites included in the OFC population were recorded from two monkeys (N and V), whereas most sites included in the MFC population were recorded from one monkey (N).

This analysis showed that LFPs in both OFC and MFC represented which option monkeys would choose on a trial-by-trial basis (Fig. 3A, B). This encoding was evident during both S1 and S2 presentations, typically at the time of the late ERP components previously described as associated with stimulus-reward values (see for comparison Fig. 2A, B). The different stimulus pairs presented on each trial also affected the performance of the classifiers trained on the OFC and MFC data. Notably, the classifiers’ choice prediction evolved as the stimuli were sequentially presented and the predicted choice often matched the actual choice, at least after both stimuli were presented (Fig. 3B).

To better understand the dynamics of these signals, we looked at how monkeys used the learned statistics of the task to augment their decisions. At the time of S1, monkeys didn’t have any information about the upcoming S2 value, but given that monkeys had significant experience with the task, it is likely that they were using the value of S1 to predict S2. This is possible in the present task because the uncertainty about the upcoming S2 value varied with the value of S1. For example, if the value of S1 was 0.2 ml on a given trial, there was a 50% chance that S2 value will be greater or lower (i.e. the maximum uncertainty in this task). On the other hand, if the value of S1 was 0 or 0.8 ml, there was no uncertainty about the following choice. Alternatively, if S1 was a stimulus associated with 0.1 or 0.4 ml this was associated with an intermediate level of uncertainty, respectively a 66.6% or 33.3% chance that S2 value will be greater than S1 (for full information on pairs, see Methods).

Taking the choice uncertainty into account we observed that the decoding performance of the decision from OFC population activity during the S1 period was strongly correlated with the estimated optimal choice probability at that time (Fig. 3C). A similar relationship was observed for the MFC population. These analyses therefore reveal that monkeys were biasing their potential choices on each trial based on the value of the first stimulus that was presented. Notably, the fact that we were able to significantly decode monkeys’ choices during S1 does not imply that the coding accurately predicted the subsequent choice. Although, knowledge about the task structure might be an efficient strategy to maximize reward, by reducing the time before reward, it might result in an incorrect prediction and a planned motor response associated with a lower value option. This planned response would need to be updated if a higher value option is presented second. Indeed, careful inspection of Figure 3A shows that strong encoding of S1 choice was reversed when the less likely S2 stimulus were presented, violating monkeys’ expectation (notably when S1/S2=0.1/0 ml and S1/S2=0.4/0.8 ml, see also Fig. 3B).

Following the presentation of S2, both OFC and MFC classifiers reached better overall performance in predicting monkeys’ actual choices (Fig. 3A, B). However, differences were observed between the two areas. In particular, the decoding accuracy from the OFC population appeared linearly scaled with the stimulus value difference (Fig. 3D). This was not the case for the MFC. Instead, the estimated probability of choosing mostly discriminated the 2 possible choices in a step-wise manner (at least when the decoding accuracy reached significance), without being affected by the difference in value between the stimuli. This is reflected by significantly higher accuracy levels in the MFC compared to the OFC in 6 different stimulus pairs (highlighted in Fig. 3D). Also, more pairs of stimuli were significantly decoded from the MFC than the OFC population after S2 presentation (time bin t2, OFC=5/14 and MFC=11/14 significantly decoded pairs; **X**^2^=5.25, p=0.02; Fig. 3B right panel).

The very low number of sites in some individual subjects (e.g. monkey H: OFC=2 and MFC=8; monkey N: MFC=3 sites) prevented us from confirming the robust existence of these effects in all subjects. As shown in Fig. 5B (top row), the choice decoding accuracy from the two remaining monkeys in OFC revealed large inter-individual variability. Ultimately, the choice classifiers could only be tested using the MFC recordings of monkey N, meaning that the effects reported should be treated with caution.

In summary, these analyses demonstrated that the ERPs recorded in the MFC, and to a lesser extent in OFC, contained information about the impending choice (S1 or S2). Together with the results on the encoding of value by ERPs, this suggests that LFPs in OFC and MFC represent distinct but complementary information associated with choice behavior.

### Amygdala lesions altered stimulus value coding

Following the acquisition of the preoperative recordings, all 3 monkeys received bilateral excitotoxic lesions of the amygdala, covering both centromedial and basolateral nuclei (Fig. 1B). Details regarding the method and extent of the lesions can be found in Rudebeck et al. (2013). We then recorded LFP signals from 298 and 184 sites postoperatively in the OFC and MFC respectively (monkey N: 169 OFC and 114 MFC; monkey H: 70 MFC; monkey V: 129 OFC). Postoperatively, the presentation of S1 and S2 elicited both the early negativity and late positivity observed before the lesion (Fig. 4A). S1 elicited an early negativity around 250 ms (average ± std, OFC=268.8±31 ms; MFC=268.8±35 ms) and a late positivity after 400 ms (OFC=425.7±69 ms; MFC=424.6±64 ms). Both components were also observed after S2 (negativity: OFC=260.2±24 ms, MFC=243.9±32 ms; positivity: OFC= 446.6±59 ms; MFC=431.5±63 ms). Amygdala lesions abolished the latency differences previously observed between the OFC and the MFC for S1 components (KW test OFC vs. MFC, negativity: H=0.02, p=0.88; positivity: H=6.3e-4, p=0.98) but not for S2 (negativity: H=27.89, p=1.28e-7; positivity: H=3.32, p=0.07). Compared to the preoperative recordings, the latency of the different components was decreased following the amygdala lesion in OFC (KW test PreOp vs. PostOp, S1 negativity: H=4.29, p=0.038; S1 positivity: H=7.34, p=0.007; S2 negativity: H=11.68, p=6.3e-4; S2 positivity: H=22.63, p=1.9e-6). This was not the case in the MFC, except for the S2 negativity (KW test PreOp vs. PostOp, S1 negativity: H=1.23, p=0.27; S1 positivity: H=1.74, p=0.187; S2 negativity: H=17.88, p=2.35e-5; S2 positivity: H=0.35, p=0.55). Finally, we also observed a clear overall decrease in the amplitude of the ERP responses in both OFC and MFC relative to the preoperative recordings (Fig. 4A).

**Figure 4.**
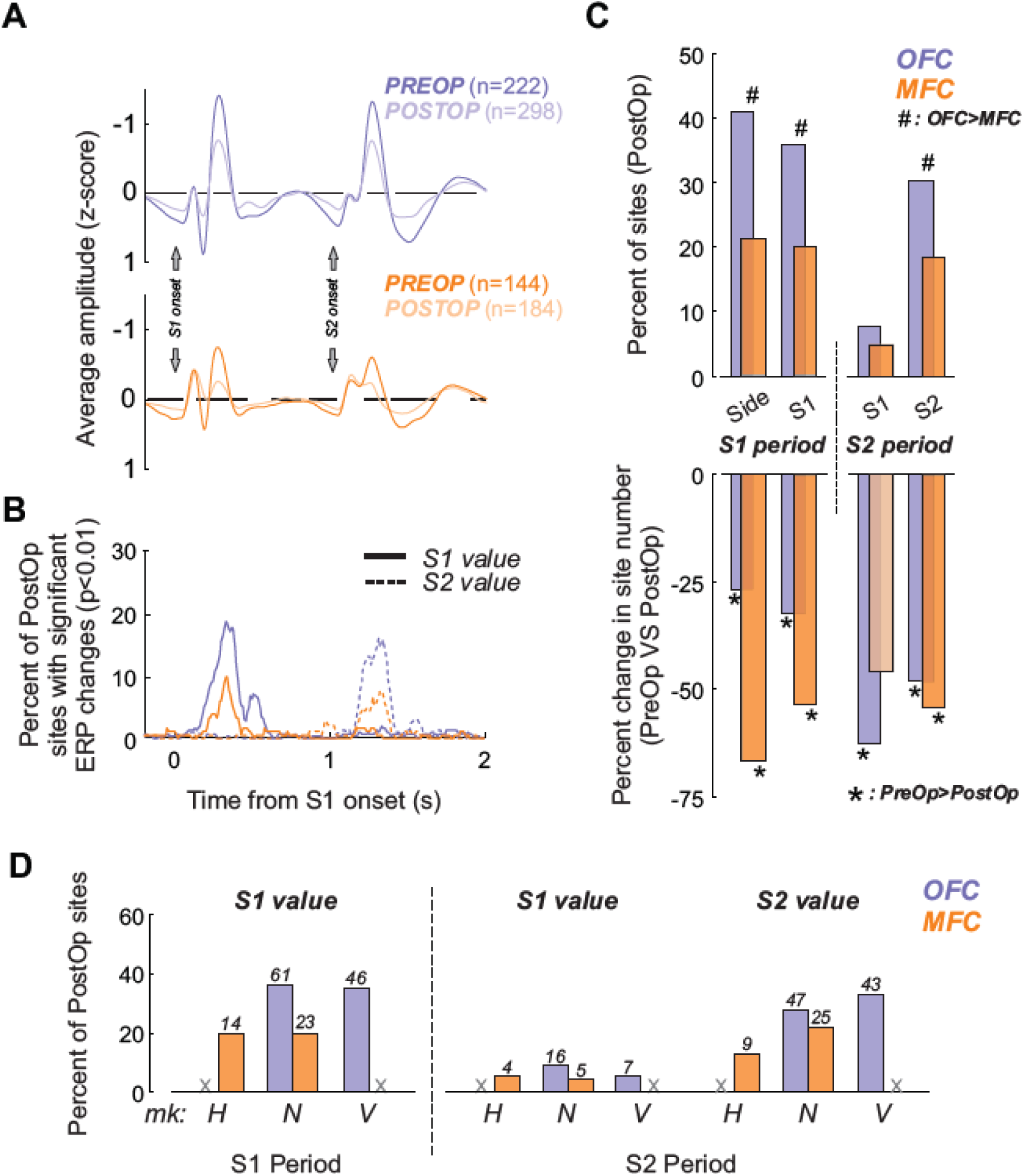
Effect of amygdala lesions on ERP signals. (**a**) Grand average normalized ERP responses induced by the presentation of S1 and S2 in the OFC (top panel) and MFC (bottom panel) before and after amygdala lesions. Black lines indicate significant differences in power between recordings acquired before and after amygdala lesions (KW test, p<0.01). (**b**) Time-resolved percentage of significant sites (hierarchical ANOVA thresholded at p<0.01, see Methods) encoding either S1 (solid lines) or S2 values (dashed lines) in both OFC and MFC. (**c**) Top panel: Percentage of sites significantly encoding stimulus side and values in the hierarchical ANOVA during S1 or S2 periods (S1: 0 to 1s; S2: 1 to 2 s) after amygdala lesion. Bottom panel: Percent change in site number compared to the preoperative data. Hashtags and stars indicate a significant difference between OFC and MFC or between PreOp and PostOp, respectively (Chi-square tests, p<0.05). (**d**) Individual monkeys’ percentage and number of sites in the OFC and MFC showing a significant effect of S1 and S2 values either during S1 or S2 periods. Grey crosses show that no recordings were performed on the given region of the corresponding monkey (H, N and V represent the three monkeys).

During the S1 period, the encoding of S1 values was still observed at a substantial number of OFC and MFC sites (OFC=107/298, 35.9%; MFC=37/184, 20.1%), with more sites in the OFC (**X**^2^=13.55, p=2.32e-4) (Fig. 4B,C). Similarly, more sites in the OFC compared to the MFC encoded S2 values during S2 period (OFC=90/298, 30.2%; MFC=34/184, 18.5%; **X**^2^=8.18, p=0.0042). This difference in proportions of site encoding S1 or S2 value was highly consistent between monkeys (Fig. 4D). More importantly, these proportions were smaller than before the amygdala lesion, for both S1 value during S1 period (PreOp vs. PostOp, OFC: **X**^2^=15.42, p=8.6e-5; MFC: **X**^2^=20.18, p=7e-6) and S2 value during S2 period (PreOp vs. PostOp, OFC: **X**^2^=40.6, p=1.8e-10; MFC: **X**^2^=19, p=1.3e-5). This decrease was robust for OFC sites in the two monkeys during both S1 (PreOp/PostOp, monkey N: 69.1/36.1%; monkey V: 44.6/35.6%) and S2 values (monkey N: 65.4/27.8%; monkey V: 54.5/33.3%). For the MFC, however, only monkey N showed a decrease in the proportion of sites encoding S1 (PreOp vs. PostOp, monkey N: 61/20.1%; monkey H: 14.3/20%) and S2 reward-values (monkey N: 58.4/21.9%; monkey H: 12.5/12.8%). Thus, caution needs to be taken when interpreting the effect of amygdala lesions on the encoding of stimulus-reward value in the MFC, given the variability between monkeys.

Furthermore, only ∼5% of sites encoded the value of S1 during the S2 period (OFC=7.7% and MCC=4.89%, **X**^2^=1.46, p=0.2259). This was lower than pre-operatively in the OFC (PreOp vs. PostOp, **X**^2^=18.69, p=1.5e-5), and was evident in both monkeys (PreOp/PostOp, monkey N: 37/9.4%; monkey V: 11.5/5.4%) (Fig. 4).

Thus, stimulus-reward value encoding by LFPs was reduced following amygdalectomy in both OFC and MFC, but differences in encoding between the areas was maintain (OFC > MFC encoding of S1 and S2). This pattern of effects is different to the changes we observed in single neuron encoding of stimulus values, where lesions reduced the difference between OFC and MFC as a result of diminished encoding in OFC (Rudebeck et al., 2013). These analyses suggest that amygdala input has different effects on single neuron and LFPs in prefrontal cortex.

Postoperatively, stimulus side was encoded at more sites in OFC than in MFC, both during S1 presentation (OFC=40.9% and MFC=21.2%, **X**^2^=19.93, p=8.01e-6) and S2 presentation (OFC=40.2% and MFC=23.3%, **X**^2^=14.51, p=1.39e-4) (Fig. 4C). Compared to before amygdala lesions, fewer sites encoded the side of the stimuli in the OFC and MFC, during both S1 (PreOp vs. PostOp, OFC: **X**^2^=11.35, p=7.5e-4; MFC: **X**^2^=61.4, p=4.7e-15) and S2 (PreOp vs. PostOp, OFC: **X**^2^=19.6, p=9.3e-6; MFC: **X**^2^=16.32, p=5.3e-5). Although the proportions were different between monkeys (as reported preoperatively), the decrease in coding was consistent across monkeys in the OFC for both S1 (PreOp/PostOp: monkey N=93.8/62.7%, monkey V=32.2/12.4%) and S2 (monkey N=92.6/63.3%, monkey V=43/10.1%). This was also true in the MFC for both S1 (monkey H=37.5/21.4%, monkey N=92.2/21.1%) and S2 (monkey H=21.4/11.4%, monkey N=64.9/30.7%).

Finally, we also found a significant decrease in the encoding of the identity of S1 or S2, either color or shape stimuli, in the OFC (S1=16/298, 5.4%; S2=21/298, 7%; PreOp vs. PostOp: **X**^2^>9.19, p<0.0024). This coding was still relatively absent in the MFC following the amygdala lesion (S1=5/184, 2.7%; S2=3/184, 1.6%; PreOp vs. PostOp: **X**^2^<2.85, p>0.09). Changes following amygdala lesions were consistent across monkeys in the OFC for both S1 (PreOp/PostOp: monkey N=12.3/3.6%, monkey V=17.4/7.8%) and S2 (monkey N=18.5/7.1%, monkey V=17.4/7.8%). This was also true in the MFC for both S1 (PreOp/PostOp: monkey H=5.4/4.3%, monkey N=6.5/1.8%) and S2 (monkey H=1.8/2.9%, monkey N=7.8/0.9%).

To summarize, we observed major alterations of the ERPs in both OFC and MFC following amygdala lesions. Although a significant proportion of sites still encoded the reward-value associated with the different stimuli, amygdalectomy markedly reduced the encoding of this aspect of the task in both areas.

### Amygdala lesions abolished the encoding of choices

We applied the same multiple linear regressions method on the postoperative population ERP activity to investigate whether the encoding of choice was affected by the removal of the amygdala. Classifiers were trained and tested on 47 randomly selected sites (total number of available sites exceeding trial number requirements, n=120 OFC and n=86 MFC). Postoperatively, it was not possible to decode monkeys’ choices using either OFC or MFC population ERP activity (Fig. 5A). The accuracy of the classifiers to predict monkeys’ choices almost never reached significance in the time-resolved analysis. Similarly, we were only able to show a significant decoding during time bin *t2* in 3/14 and 2/14 pairs in OFC and MFC, respectively (see white stars in Fig. 5A, bottom panel). Consistent results were observed in the three subjects (Fig. 5B). This reveals the major influence of the amygdala in the computation of choice-related activity in the MFC. Despite this change in MFC, it is important to keep in mind that monkeys were still able to perform the task, with similar near-optimal performance than before lesions.

**Figure 5.**
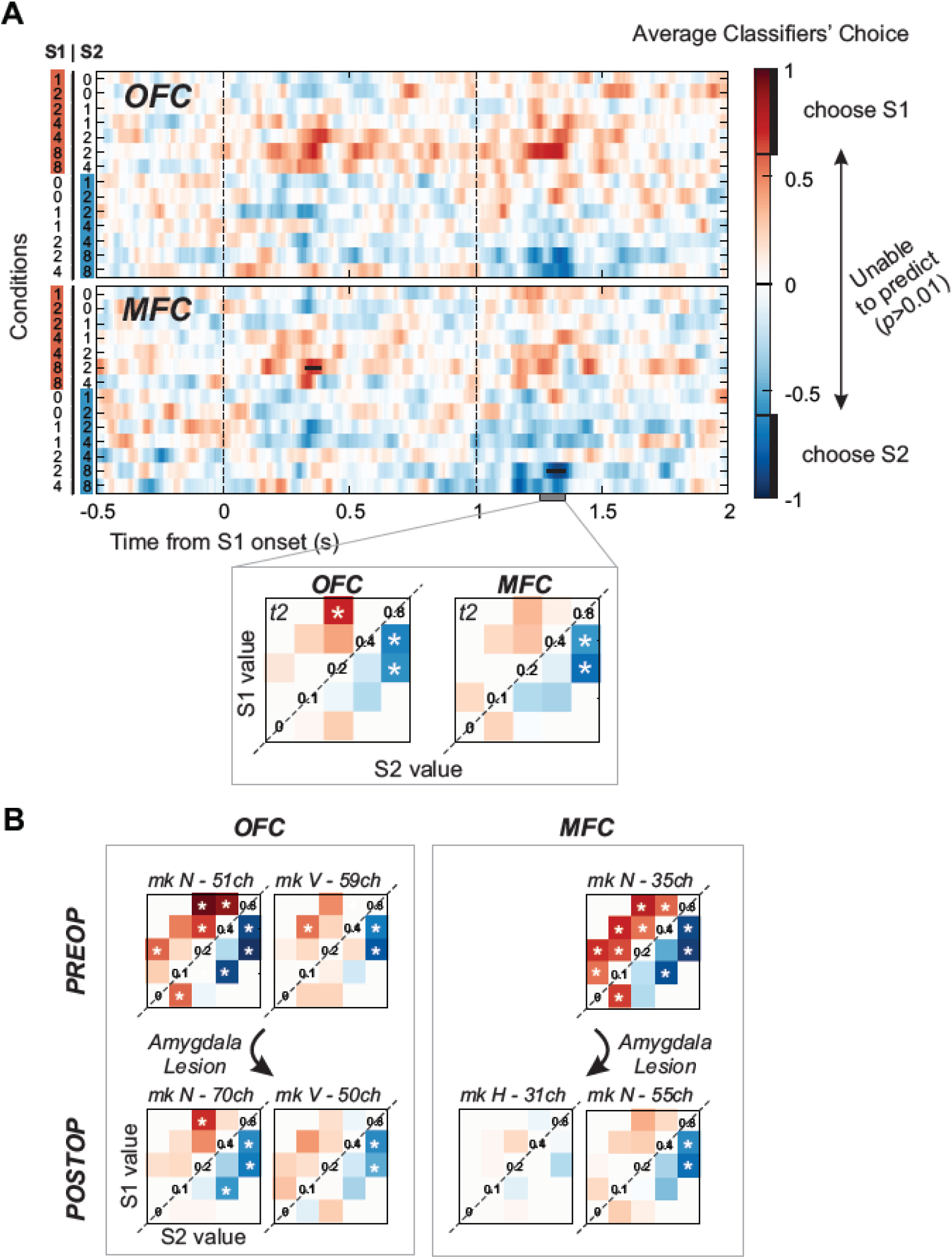
Amygdala lesions abolished the encoding of choices in the ERP. (**a**) Average classifiers’ choice performance after amygdala lesion (red=S1 and blue=S2) for each individual pairs of stimuli (labeled as *Conditions*, y-axis) over time (x-axis). Black bars represents a significant statistical preference for choosing S1 or S2 based on permutation testing (threshold at p<0.01 for at least 6 consecutive time bins). Inset at the bottom represents the average choice performance during time bin t2. (**b**) Average classifiers’ choice performance derived from individual monkeys during time bin *t2* for each S1/S2 pairs and for both OFC (left) and MFC (right) populations, before and after amygdala lesions. White stars represent a significant performance for the considered S1/S2 pair (p<0.01).

### ERPs and single neuron activity convey different information related to choices

Preoperatively, we found that choice decoding of ERP signals was more accurate in the MFC compared to OFC (Figs 3B,C). Further, our analyses showed that amygdala lesions almost abolished the encoding differences between OFC and MFC and the ability to decode the monkey’s choices. By contrast, individual neurons in both OFC and MFC are only weakly tuned to monkeys’ choices during this task (Rudebeck et al. 2013), revealing a possible dissociation between the information carried by single neurons and LFPs. However, different methods were applied with the two datasets, and it is possible that using population decoding measures on the single neuronal recordings might reveal other aspects of choice-related signaling. We therefore applied the same decoding analyses to both measures of neural activity (see Methods). As before, the number of predictors (ERP sites or neurons) was similar to avoid potential nonspecific biases of classifiers’ performance.

It was possible to decode monkeys’ choice using either single neuron or ERP activity recorded in OFC or MFC, at least when specific S1/S2 pairs where presented (shown for time bin t2 in Fig. 6A). Overall, classifiers using single neuron activity from either OFC or MFC reached similar decoding performance to that from ERPs recorded in OFC, although the significantly decoded pairs differed between the two measures and brain regions. Following amygdala lesions, we observed a decrease in the classifier’s accuracy compared to the preoperative single neurons recordings; we were unable to decode monkeys’ choices in as many S1/S2 pairs (Fig. 6B, bar plot). This occurred in both MFC and OFC. Interestingly, lesions were not simply associated with decreased classifier performance; a significant increase in performance was observed in a few S1/S2 pairs (4 and 2 pairs for OFC and MFC respectively; see upward arrows in Fig. 6A, bottom panel). This analysis indicates that the information contained in single neuron and ERP populations was differentially affected by amygdalectomy.

**Figure 6.**
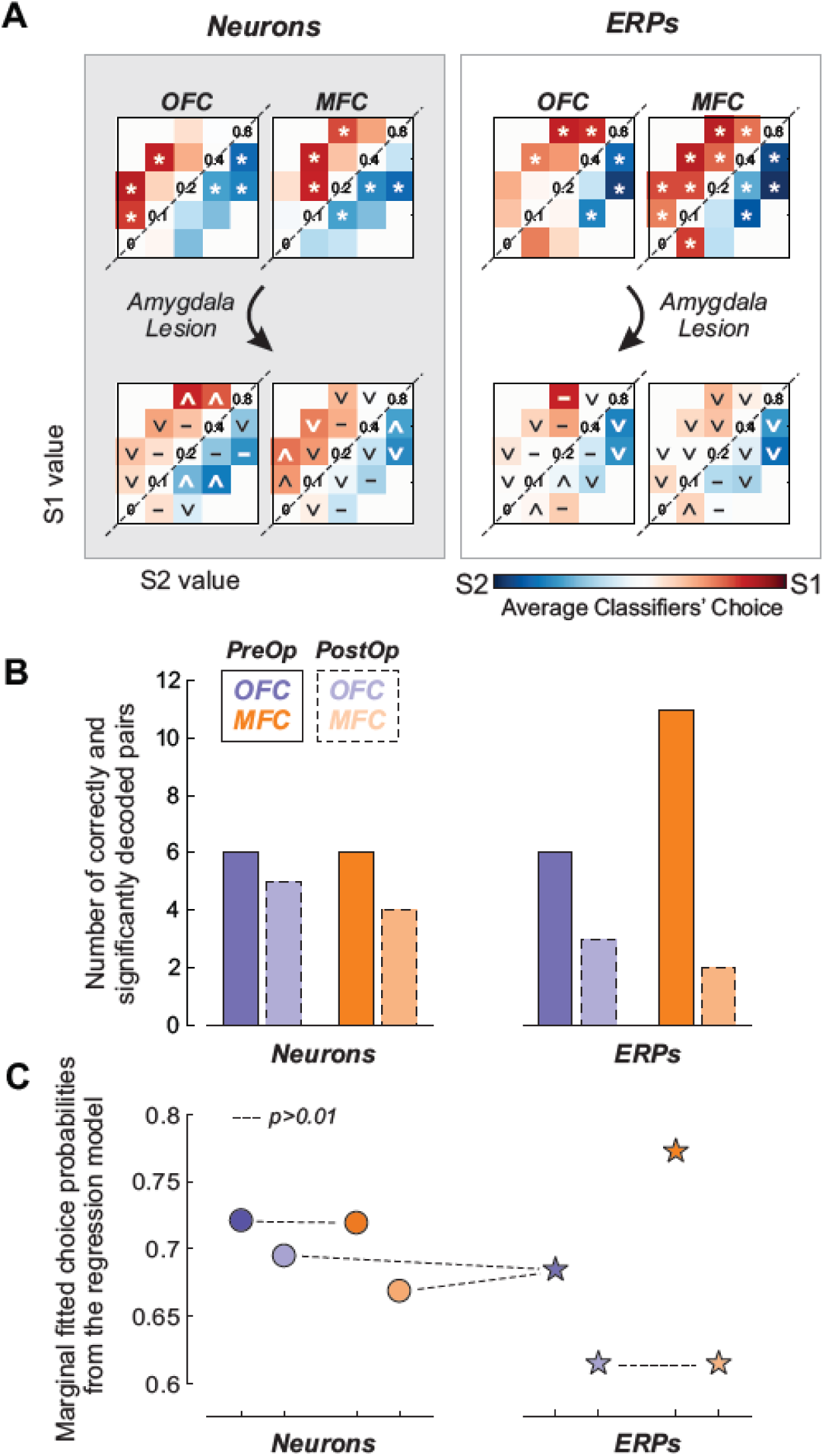
Single neurons and ERPs differences in the encoding of choices. (**a**) Average classifiers’ choice performance during time bin *t2* for each S1/S2 pairs and for both single neuron (left) and ERP (right) populations. White marks indicate a significant decoding performance for the considered S1/S2 pair (stars in PreOp, increase / null / decrease marks in PostOp). Marks for PostOp represent significant (increase or decrease signs) and non-significant changes (dash) when compared to the PreOp populations. (**b**) Number of significantly and accurately decoded S1/S2 pairs for the different population considered (**c**) Overall choice probabilities extracted from the mixed-effect logistic regression for each population. Non-significant post-hoc comparisons are represented by dash lines (*p*>0.01). Conventions as in previous figures.

To statistically compare whether the overall choice coding strength was different in the different neural signals and/or modulated following amygdala lesions, we fitted a mixed-effect logistic regression to the output from the different classifiers (see Methods). Categorical fixed-effect factors included recording types (single neurons vs. LFPs), areas (OFC vs. MFC) and surgery (PreOp vs. PostOp). S1/S2 pairs were included as a random-effect factor, allowing only changes in the intercept (i.e. choosing more S1 or S2 depending on their values). The model results are summarized in Fig. 6C. All interactions survived model selection and were statistically significant (Type x Area: t_(104,1)_=11.71, p=1.03e-20; Type x Surgery: t_(104,1)_=-5.16, p=1.1e-6; Area x Surgery: t_(104,1)_=-3.25, p=1.5e-3). The three-way interaction (Type x Area x Surgery) also remained in the best model (t_(104,1)_=-6.13, p=1.6e-8), highlighting the existence of a strong dissociation between the factors considered. Post-hoc comparisons using a threshold at *p*<0.01 revealed that: 1) choices were more strongly encoded by single neurons compared to ERPs in the OFC, whereas the opposite was true for the MFC (Fig. 6C). 2) Choice encoding was also significantly greater in MFC than OFC, but only for ERPs, not single neurons (Fig. 6C). 3) Amygdala lesions significantly reduced the performance of the classifiers on both single neurons and ERPs, although the effect was more pronounced for ERPs (relative to that for single neurons) and in the MFC populations (relative to OFC). In summary, amygdala lesions differentially affected choice coding in OFC and MFC at the level of single neurons and ERPs. The most prominent change was the reduction in spike encoding of choices in OFC and the decrease ERP encoding of choices in MFC.

## DISCUSSION

Here we analyzed the local field potentials recorded in OFC and MFC of monkeys engaged in a task where they chose between two sequentially presented stimuli associated with different sized fluid rewards on each trial. Before lesions of the amygdala, ERPs in OFC, and to a lesser extent in MFC, encoded the reward value of the two stimuli presented (Fig. 2). Furthermore, if a site encoded the value of S1 it was highly likely to encode the value of S2. This closely matched findings from our previously published single neuron recordings. Despite this correspondence, we found that there was no direct relationship between the encoding of value by single neurons and ERPs simultaneously recorded on the same electrode. We then looked at how ERPs encoded the choices that the monkeys would make on each trial. We found that ERPs recorded in MFC, and to a lesser extent in OFC, contained relevant information about the upcoming choices that monkeys would make (Fig 3). Taken together, the findings indicate that local field potentials in OFC and MFC represent the relevant information to make adaptive and optimal decisions.

Removing amygdala input to OFC and MFC strongly reduced ERP encoding of stimulus reward value in both areas (Fig. 4). It also decreased the encoding of monkey’s choices. This was most apparent in MFC where the lesions completely abolished choice-related signals encoding (Fig. 5). When we compared the effects of lesions on ERP and single neuron encoding of choices, the lesions appeared to mostly affect ERP, not single neuron. This was especially prominent in MFC (Fig. 6). Taken together these data suggest that amygdala inputs are important for augmenting reward-value and choice signals in PFC, most notably ERP choice-related signals in MFC.

### Encoding of reward-value and choices in OFC and MFC

Both OFC and MFC have been linked to reinforcement-guided decision making, notably when monkeys have to choose between different stimuli or courses of action associated with reward (e.g., Thorpe et al., 1983; Matsumoto et al., 2003; Wallis and Miller, 2003; Amiez et al., 2006; Kennerley et al., 2006; Walton et al., 2010; Stoll et al., 2016). It has also been emphasized that encoding in OFC and MFC is not identical (Kennerley et al., 2009, 2011), and that each area makes distinct contributions to different aspects of decision-making (Rudebeck et al., 2008; Camille et al., 2011). Here, we observed that both brain structures appeared to reflect stimulus-reward values and the upcoming choices. However, the strength of coding of each factor differed between OFC and MFC and a clear dissociation was apparent: ERPs in OFC strongly encoded the reward value associated with the stimuli presented on each trial whereas ERPs in MFC were more closely aligned to the *product* of value signals, reflecting the encoding of monkeys’ choices. The effect in MFC should, however, be taken with caution as decoding choice-related signals was performed using a non-homogeneous and limited number of channels from the different monkeys (see Fig. 5B). Nevertheless, this could be related to our previous observation, where single neuron encoding of the amount of reward associated with S2 in MFC was more influenced by the value of S1 (i.e., was more akin to a relative valuation; Rudebeck et al., 2013).

Indeed, based on work in humans, encoding of choice in MFC might be expected. MFC and medial OFC have both been proposed to play a role in the comparison process depending on the context (Rushworth et al., 2012). In addition, MFC is critical for combining multiple variables important for the decision processes, including both costs and benefits (Rudebeck et al., 2006; Kennerley et al., 2009; Stoll et al., 2016). Hence, it has been argued that the MFC could represent the value of exploring alternative courses of actions (Kolling et al., 2016). Although our task doesn’t specifically depend on action values, representing which actions have been performed and their potential value could be a crucial part of deciding whether to stay engaged in the task.

It is important to note that the design of our task and highly consistent choice patterns of our subjects prevented us from fully disentangling value and choice-related signals (as is the case in nearly every value-based decision-making task). As noted earlier, this aspect of the design was necessary to ensure that any alterations in neural activity consequent to the lesion could be interpreted. Because of this aspect of our study, monkeys’ choice information could be seen as a binary version of the difference in value of both stimuli. Therefore, the dissociation we observed between OFC and MFC could be linked to the specific way value-related information is represented in these regions.

Nevertheless, both OFC and MFC ERPs contained information about stimulus-reward values and choices. This observation supports the notion of permeability of information throughout the PFC. This could be the result of both the anatomical and functional properties of the PFC. First, strong anatomical connections exist between the different parts of PFC, notably between the OFC and the MFC (Carmichael and Price, 1996). Also, the associative role that has been attributed to PFC, as well as the broad influence of motivational factors on this structure, makes it suitable to represent multiple parameters related to value and decisions (Wallis and Rich, 2011). Our results support the view that OFC and MFC, albeit being tuned differently by value and choice information, work in unison when deciding between alternatives in an adaptive and optimal manner.

### Correspondence between ERPs and single neuron activity

Despite the potential for LFPs to shed light on information processed in PFC, only a handful of studies have assessed value and/or choice encoding by LFPs in PFC (Morrison et al., 2011; Hunt et al., 2015; Rich and Wallis, 2016). We found that value encoding in ERP signals was not predictive of whether a neuron recorded at the same location would encode value as well. This result is somewhat surprising as it suggests that there are differences in the type of information carried in these two measures of neural activity, and, ultimately, that they are not simply the same process looked at from different angles. One simple explanation for this dissociation is that differences between the two measures (continuous signal vs point process) mean that it is difficult to truly compare the signals. This could be an especially acute problem in OFC where spiking activity is typically sparse (e.g. Thorpe et al., 1983). Against this, however, there have been reports of differences in the type of information signaled between LFPs and single neuron activity in parts of temporal cortex (Kreiman et al., 2006; Nielsen et al., 2006) and PFC in particular (Monosov et al., 2008; Lara and Wallis, 2014; but see Rich and Wallis, 2016), indicating that this may be a valid difference.

An alternative explanation relates to the basis of ERP signals. ERPs are commonly assumed to reflect postsynaptic events of thousands of neurons, depending on the recording setup. Therefore, ERPs are thought to partly represent the input to a region from other cortical or sub-cortical regions (Nguyen and Lin, 2014). However, given that many synapses between neurons are short-range (i.e., within-area), ERPs could contain a mixture of information from both inputs and outputs of a region (Douglas and Martin, 2004); by contrast, single neuron activity only reflects the output of a region.

While there was a location specific dissociation between single neuron and ERP encoding of value, at the population level both measures were highly similar (compare example Fig. 2C with Fig. 3 in Rudebeck et al., 2013). This close correspondence, however, was absent for one of our findings: encoding of choice by ERPs in MFC (Fig. 6). Given the basis of the ERP signal noted above, it is possible that MFC receives choice-related signals from amygdala (Fig. 6) or potentially other parts of PFC that require amygdala input, but this does not result in a strong cascade of activity in individual neurons. Apart from amygdala, such choice signal could potentially come from medial OFC where comparison related activity has been reported in both macaques and humans (Boorman et al., 2009; Kolling et al., 2012; Strait et al., 2014). The lack of single neuron encoding of choices during the present task could be linked to the relatively low involvement of the MFC and OFC in this task, where values were already learned. By contrast, if the value of stimuli changed unexpectedly or has to be learned, this might trigger a cascade of events throughout the PFC, increasing the need for cognitive control which could potentiate the processing of choice and value information in the MFC and OFC, respectively. This idea would appear to fit with data from the same subjects showing that during stimulus-reward learning, single neuron activity in MFC closely matches stimulus values and is indistinguishable from OFC encoding (Rudebeck et al., 2017).

### Amygdala influence on reward value and choice coding

We found that amygdala lesions not only strongly affected ERP correlates of stimulus-reward values and choices, but also the encoding of other parameters such as the stimulus side or identity (Fig. 4). This is different to the effect of amygdala lesions on the activity of individual neurons, where there was not a significant reduction in the encoding of these parameters (Rudebeck et al., 2013). A wholesale reduction in ERP amplitude following lesions could be responsible for the decrease in coding in the different parameters. Yet ERP amplitudes in OFC postoperatively were still higher than MFC preoperatively, but nevertheless less tuned to the different factors considered. Therefore, such changes might represent a specific loss of information as opposed to a nonspecific reduction in signal-to-noise ratio. In fact, neurons in the amygdala have been shown to reflect both the identity and the location of a stimulus, possibly related to the role of amygdala in directing attentional processes (Peck et al., 2013, 2014).

Previous studies where functional measures, either single neuron recording or functional MRI, have been combined with amygdala lesions during reward based tasks have reported changes in encoding in PFC (Schoenbaum et al., 2003; Hampton et al., 2007). Although our findings are in broad agreement with this previous work in humans and rodents, the change in choice encoding in MFC that we found is most pertinent to a human fMRI study detailing the effects of amygdala damage (Hampton et al., 2007). Hampton and colleagues reported that expected reward signals in MFC (i.e., the outcome of decisions) were reduced in two humans with amygdala damage performing a reward-guided task. Our data therefore confirm and extend these findings by showing the dynamic nature of these effects and that choice signals are abolished irrespective of when the stimuli are presented (decoding on S1 or S2, Fig. 5).

Given the strong reciprocal projections between PFC and amygdala (Morecraft and Van Hoesen, 1998; Ghashghaei et al., 2007), it could be argued that the changes we observed were the result of the loss of amygdala inputs to the OFC and MFC. This could be supported by the relatively greater decrease in the encoding of value and choice in ERPs than in single neurons given that ERPs could reflect inputs to these regions. However, we cannot rule out the possible influence of other brain regions. For example, the loss of information related to value and choice in the MFC could be an indirect result of the loss of amygdala inputs to the medial and lateral OFC, which receive strong projections from the amygdala (Ghashghaei et al., 2007). Alternatively, thalamic and dopaminergic inputs could play a role in the transmission of information from the amygdala to the PFC (Williams and Goldman-Rakic, 1998; Timbie and Barbas, 2015).

### Summary

We recorded ERPs in the OFC and MFC of macaque monkeys while they performed a reward-based choice task. We found that OFC and MFC carry distinct signals related to decision-making at the level of single neurons and LFPs. While both single neurons and LFPs in OFC predominantly encoded stimulus-reward values, ERP signals in MFC were specifically related to monkey’s choices. This correlate of decision-making in MFC was unique in that it was not seen in the activity of single neurons and it was almost entirely dependent on input from the amygdala. Given the prominent role of MFC-amygdala interactions in numerous psychiatric disorders, alterations in choice-related signals in MFC could be used as a marker of amygdala dysfunction.

## ACKNOWLEDGEMENTS

This work was supported by a National Institute of Mental Health BRAINS award to PHR (R01 MH110822), a young investigator grant from the Brain and Behavior Foundation (NARSAD) to PHR, a Philippe Foundation award to FMS, seed funds from the Icahn School of Medicine at Mount Sinai to PHR, and the Intramural Research Program of the NIMH. We thank Kevin Blomstrom, Kevin Fomalont and Joshua Ripple for assistance with data collection, and James Fellows, Ping Yu Chen and David Yu for help with surgery and histology.

## REFERENCES

Almeida JRC de, Versace A, Mechelli A, Hassel S, Quevedo K, Kupfer DJ, Phillips ML (2009) Abnormal Amygdala-Prefrontal Effective Connectivity to Happy Faces Differentiates Bipolar from Major Depression. Biol Psychiatry 66:451–459.

Amiez C, Joseph JP, Procyk E (2006) Reward encoding in the monkey anterior cingulate cortex. Cereb cortex 16:1040–1055.

Astrand E, Enel P, Ibos G, Dominey PF, Baraduc P, Ben Hamed S (2014) Comparison of classifiers for decoding sensory and cognitive information from prefrontal neuronal populations. PLoS One 9.

Baxter MG, Parker A, Lindner CCC, Izquierdo AD, Murray E a (2000) Control of response selection by reinforcer value requires interaction of amygdala and orbital prefrontal cortex. J Neurosci 20:4311–4319.

Boorman ED, Behrens TEJ, Woolrich MW, Rushworth MFS (2009) How Green Is the Grass on the Other Side? Frontopolar Cortex and the Evidence in Favor of Alternative Courses of Action. Neuron 62:733–743.

Camille N, Tsuchida A, Fellows LK (2011) Double Dissociation of Stimulus-Value and Action-Value Learning in Humans with Orbitofrontal or Anterior Cingulate Cortex Damage. J Neurosci 31:15048–15052.

Carmichael ST, Price JL (1996) Connectional networks within the orbital and medial prefrontal cortex of macaque monkeys. J Comp Neurol 371:179–207.

Douglas RJ, Martin K a C (2004) Neuronal circuits of the neocortex. Annu Rev Neurosci 27:419–451.

Dutta A, McKie S, Deakin JFW (2014) Resting state networks in major depressive disorder. Psychiatry Res 224:139–151.

Einevoll GT, Kayser C, Logothetis NK, Panzeri S (2013) Modelling and analysis of local field potentials for studying the function of cortical circuits. Nat Rev Neurosci 14:770–785.

Felix-Ortiz AC, Burgos-Robles A, Bhagat ND, Leppla CA, Tye KM (2016) Bidirectional modulation of anxiety-related and social behaviors by amygdala projections to the medial prefrontal cortex. Neuroscience 321:197–209.

Floresco SB, Ghods-Sharifi S (2007) Amygdala-prefrontal cortical circuitry regulates effort-based decision making. Cereb Cortex 17:251–260.

Ghashghaei HT, Hilgetag CC, Barbas H (2007) Sequence of information processing for emotions based on the anatomic dialogue between prefrontal cortex and amygdala. Neuroimage 34:905–923.

Hampton AN, Adolphs R, Tyszka MJ, O’Doherty JP (2007) Contributions of the Amygdala to Reward Expectancy and Choice Signals in Human Prefrontal Cortex. Neuron 55:545–555.

Hunt LT, Behrens TEJ, Hosokawa T, Wallis JD, Kennerley SW (2015) Capturing the temporal evolution of choice across prefrontal cortex. Elife 4:e11945.

Izquierdo A, Murray E a (2007) Selective bilateral amygdala lesions in rhesus monkeys fail to disrupt object reversal learning. J Neurosci 27:1054–1062.

Kennerley SW, Behrens TEJ, Wallis JD (2011) Double dissociation of value computations in orbitofrontal and anterior cingulate neurons. Nat Neurosci 14:1581–1589.

Kennerley SW, Dahmubed AF, Lara AH, Wallis JD (2009) Neurons in the frontal lobe encode the value of multiple decision variables. J Cogn Neurosci 21:1162–1178.

Kennerley SW, Walton ME, Behrens TEJ, Buckley MJ, Rushworth MFS (2006) Optimal decision making and the anterior cingulate cortex. Nat Neurosci 9:940–947.

Kolling N, Behrens TEJ, Mars RB, Rushworth MFS (2012) Neural mechanisms of foraging. Science 336:95–98.

Kolling N, Behrens TEJ, Wittmann MK, Rushworth MFS (2016) Multiple signals in anterior cingulate cortex. Curr Opin Neurobiol 37:36–43.

Kreiman G, Hung CP, Kraskov A, Quiroga RQ, Poggio T, DiCarlo JJ (2006) Object selectivity of local field potentials and spikes in the macaue inferior temporal cortex. Neuron 49:433–445.

Lara AH, Wallis JD (2014) Executive control processes underlying multi-item working memory. Nat Neurosci 17:876–883.

Matsumoto K, Suzuki W, Tanaka K (2003) Neuronal correlates of goal-based motor selection in the prefrontal cortex. Science 301:229–232.

Monosov IE, Trageser JC, Thompson KG (2008) Measurements of Simultaneously Recorded Spiking Activity and Local Field Potentials Suggest that Spatial Selection Emerges in the Frontal Eye Field. Neuron 57:614–625.

Morecraft RJ, Van Hoesen GW (1998) Convergence of limbic input to the cingulate motor cortex in the rhesus monkey. Brain Res Bull 45:209–232.

Morrison SE, Saez A, Lau B, Salzman CD (2011) Different Time Courses for Learning-Related Changes in Amygdala and Orbitofrontal Cortex. Neuron 71:1127–1140.

Nguyen DP, Lin SC (2014) A frontal cortex event-related potential driven by the basal forebrain. Elife 3:e02148.

Nielsen KJ, Logothetis NK, Rainer G (2006) Dissociation between local field potentials and spiking activity in macaque inferior temporal cortex reveals diagnosticity-based encoding of complex objects. J Neurosci 26:9639–9645.

Oostenveld R, Fries P, Maris E, Schoffelen JM (2011) FieldTrip: Open source software for advanced analysis of MEG, EEG, and invasive electrophysiological data. Comput Intell Neurosci 2011:156869.

Peck CJ, Lau B, Salzman CD (2013) The primate amygdala combines information about space and value. Nat Neurosci 16:340–348.

Peck EL, Peck CJ, Salzman CD (2014) Task-Dependent Spatial Selectivity in the Primate Amygdala. J Neurosci 34:16220–16233.

Pezawas L, Meyer-Lindenberg A, Drabant EM, Verchinski BA, Munoz KE, Kolachana BS, Egan MF, Mattay VS, Hariri AR, Weinberger DR (2005) 5-HTTLPR polymorphism impacts human cingulate-amygdala interactions: a genetic susceptibility mechanism for depression. Nat Neurosci 8:828–834.

Rich EL, Wallis JD (2016) Decoding subjective decisions from orbitofrontal cortex. Nat Neurosci 19:973–980.

Rudebeck PH, Behrens TE, Kennerley SW, Baxter MG, Buckley MJ, Walton ME, Rushworth MFS (2008) Frontal cortex subregions play distinct roles in choices between actions and stimuli. J Neurosci 28:13775–13785.

Rudebeck PH, Mitz AR, Chacko R V., Murray EA (2013) Effects of amygdala lesions on reward-value coding in orbital and medial prefrontal cortex. Neuron 80:1519–1531.

Rudebeck PH, Ripple JA, Mitz AR, Averbeck BB, Murray EA (2017) Amygdala Contributions to Stimulus-Reward Encoding in the Macaque Medial and Orbital Frontal Cortex during Learning. J Neurosci 37:2186–2202.

Rudebeck PH, Walton ME, Smyth AN, Bannerman DM, Rushworth MFS (2006) Separate neural pathways process different decision costs. Nat Neurosci 9:1161–1168.

Rushworth MFS, Kolling N, Sallet J, Mars RB (2012) Valuation and decision-making in frontal cortex: one or many serial or parallel systems? Curr Opin Neurobiol 22:946–955.

Schoenbaum G, Setlow B, Saddoris MP, Gallagher M (2003) Encoding predicted outcome and acquired value in orbitofrontal cortex during cue sampling depends upon input from basolateral amygdala. Neuron 39:855–867.

Stoll FM, Fontanier V, Procyk E (2016) Specific frontal neural dynamics contribute to decisions to check. Nat Commun 7:11990.

Strait CE, Blanchard TC, Hayden BY (2014) Reward value comparison via mutual inhibition in ventromedial prefrontal cortex. Neuron 82:1357–1366.

Thorpe SJ, Rolls ET, Maddison S (1983) The orbitofrontal cortex: Neuronal activity in the behaving monkey. Exp Brain Res 49:93–115.

Timbie C, Barbas H (2015) Pathways for Emotions: Specializations in the Amygdalar, Mediodorsal Thalamic, and Posterior Orbitofrontal Network. J Neurosci 35:11976–11987.

Wallis JD, Miller EK (2003) Neuronal activity in primate dorsolateral and orbital prefrontal cortex during performance of a reward preference task. Eur J Neurosci 18:2069–2081.

Wallis JD, Rich EL (2011) Challenges of Interpreting Frontal Neurons during Value-Based Decision-Making. Front Neurosci 5:124.

Walton ME, Behrens TEJ, Buckley MJ, Rudebeck PH, Rushworth MFS (2010) Separable Learning Systems in the Macaque Brain and the Role of Orbitofrontal Cortex in Contingent Learning. Neuron 65:927–939.

Williams SM, Goldman-Rakic PS (1998) Widespread origin of the primate mesofrontal dopamine system. Cereb Cortex 8:321–345.

